# Medial Prefrontal Cortex controlling the immobility of rats in the forced swimming test: a systematic review and meta-analysis

**DOI:** 10.1101/2021.04.27.441685

**Authors:** K. Domingues, F. F. Melleu, C. Lino de Oliveira

**Author notes:** **Corresponding Author**: Cilene Lino de Oliveira, Departamento de Ciências Fisiológicas, Centro de Ciências Biológicas, Universidade Federal de Santa Catarina, Campus Universitário Trindade, 88049-900 – Florianópolis – SC – Brazil., **Contact**.

## Abstract

The medial prefrontal cortex (mPFC) belong to the neural circuitry responsible for the behavioral responses to antidepressants in humans or animals. In the forced swimming (FST), a predictive test for antidepressants in laboratory rodents, inhibition, or stimulation of mPFC may produce antidepressant-like, unlike or no behavioral effect at all. Controversial findings may result from the variety of subregions of mPFC controlling behaviour of rats in the FST. The aim in the present study was to estimate the contribution of subregions of the mPFC to the control of rat behavior in the FST. For an unbiased view and well-powered analysis of the mentioned effects, a systematic review at Medline (Pubmed) followed by a meta-analysis was performed. Compared to other subdivisions, inhibition of prelimbic or infralimbic mPFC caused a significant drop of immobility time in the FST, which is an antidepressant-like effect. Summarizing, prelimbic or infralimbic cortices seem more relevant than other subregions to the control of immobility in the FST underlying the effects of antidepressants on mood and behaviour.

## 1. Introduction

The prefrontal cortex (PFC) and subdivisions may control cognitive and emotional functions through to afferent and efferent connections with cortical and subcortical areas. The PFC receives projections from visual and auditory cortices, intraparietal and posterior cingulate areas controlling the attentional processes, thalamic nuclei involved in working or long term memory, and also from sectors of the amygdala controlling emotional memory (Barbas, 2000; Petrides and Pandya, 2002). Regarding the afferent projections, the PFC connects with the premotor cortex and brainstem controlling head and body movements and with hypothalamic areas controlling visceral and behavioral responses to emotions (Barbas, 2000).

Patients with damaged ventromedial PFC (vmPFC) presented decision-making impairments (Bechara and Damasio, 2005; Bechara et al., 1994; Bechara et al., 2003; Bechara et al., 1997) and weak expression of emotions processed in subcortical regions, such as anxiety (Cunningham and Zelazo, 2007; Davidson, 2002; Dixon and Christoff, 2014; Haber and Behrens, 2014; Ochsner and Gross, 2005). Deficiencies in the PFC function has also been linked to the onset of affective disorders such as depression (Drevets et al., 2008; Farb et al., 2010; Greicius et al., 2007; Mayberg et al., 2005) or anxiety (Bishop et al., 2004; Chamberlain et al., 2008; Davidson, 2002; Goldin et al., 2009) and bipolar disorders (Blumberg et al., 2003; Blumberg et al., 1999; Dixon et al., 2017; Frye et al., 2007).

Treatments for affective or anxiety disorders also affect the function of PFC (Carey et al., 2004; Crane et al., 2017; Evans et al., 2009; George et al., 2000; Whalen et al., 2008). For example, depressed or anxious patients benefited from daily treatment of PFC with transcranial magnetic stimulation (TMS) for two weeks (GEORG et al., 2000). Besides, treatment with the antidepressant citalopram (a selective serotonin reuptake inhibitor, SSRI) for eight weeks resulted in deactivation of the anterior cingulate cortex (aCg) in patients with anxiety disorders, although no pattern of cortical activation predicted response to subsequent pharmacotherapy (CAREY et al., 2004). In depressed patients, functional magnetic resonance imaging (fMRI) showed a task-induced activity of vmPFC and anterior cingulate cortex (aCg) as a predictor to remission after treatment with escitalopram or reboxetine (Crane et al., 2017). Moreover, patients manifesting lower activity in vmPFC and aCg may better profit from antidepressant treatment than those with high activity in these areas (Crane et al., 2017), indicating that low activity of the PFC might be either cause or consequence of the antidepressant effect. Studies in laboratory animals may help to establish the directions of a putative cause-effect relationship between PFC and antidepressant treatment.

In animal models, PFC also seems to contribute to the control of the behavioral effects of antidepressants (Sartim et al., 2016; Slattery et al., 2011). The inhibition of medial PFC (mPFC) or infralimbic cortex (IL) in rats (Sartim et al., 2016; Slattery et al., 2011) reduced immobility time in the forced swimming test (FST), a predictive test for the action of antidepressants in laboratory rodents (Porsolt et al., 1977; Porsolt et al., 1978). Injection of the GABA_A_ agonist muscimol into mPFC reduced immobility time of rats in the FST, which is considered an antidepressant-like effect (Slattery et al., 2011). The 5-HT_1A_ agonist 8-OH-DPAT injected in the IL also produced an antidepressant-like effect (Sartim et al., 2016). However, other studies reported absence of antidepressant-like effect s by inhibiting the prelimbic (PL) or aCg or vmPFC in rats tested in the FST (Li et al., 2012; Burgdorf et al., 2013; Chang et al., 2014). These incongruent findings preclude a final appraisal on the role of PFC on antidepressant-like effects in animal models indicating different roles for different subdivisions (Figure 1).

**Legend for Figure 1.**
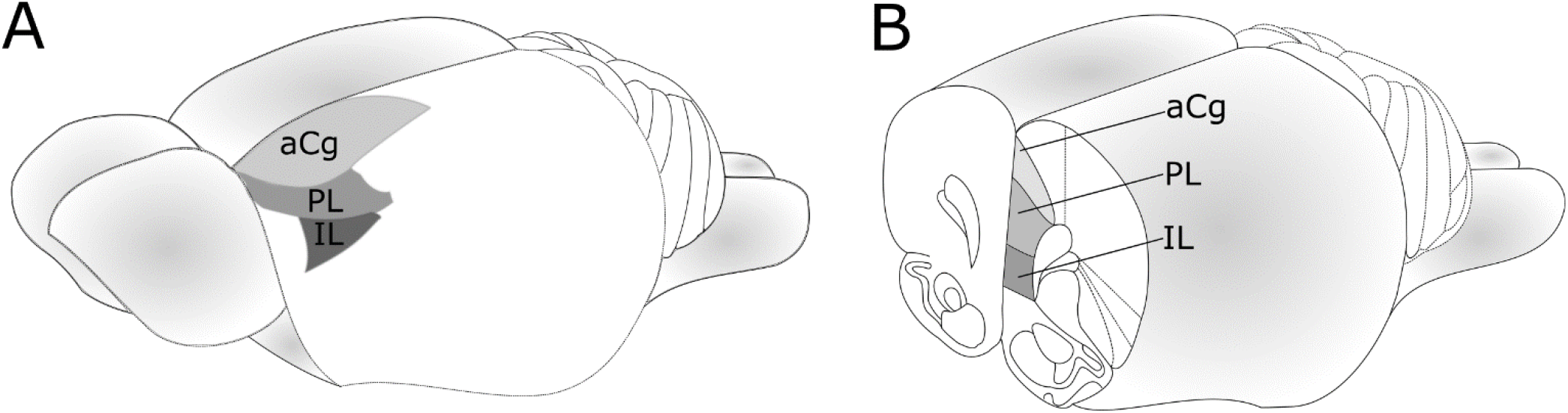
Anatomical subdivisions of the prefrontal cortex of the rat. A) Lateral view of the rat’s brain. Shaded areas are showing the PFC and its subregions. B) Coronal section through rostral telencephalon. Shaded areas showing the PFC and subregions. Abbreviations: **aCg:** Anterior Cingulate cortex, **PL:** Prelimbic cortex, **IL:** Infralimbic cortex.

The forced swimming test is a behavioral test with a predictive value, that is, a decreased immobility time may suggest that a certain compound may have an antidepressant activity.

The first session, the pre-test, is used to induce the immobility behavior, also referred to as behavioral despair, which is similar to the learned helplessness. This is the best condition to assess antidepressant effects.

It is the most used test to screen antidepressant compounds.

It is sensible to most of the drugs used as antidepressant in humans and can differentiate drugs that do not have antidepressant activity, such as benzodiazepines. Other tests that rely on spontaneous or conditioned behaviors do not express the same sensibility to differ between antidepressants and other psychoactive drugs.

However, recent discussions stablish the immobility behavior as being more adaptive in this context, being a way to conserve energy until better chances of escape are presented.

Still, the FST is a tool to investigate the neural substrate involved in antidepressant action and response.

To address the hypothesis of different contributions of discrete subregions of the PFC to the control of rat behavior in the FST, a review of the available literature was performed. For a non-biased vision of the bibliography and well-powered analysis, a systematic review of the scientific literature followed by a meta-analysis was conducted according to a pre-planned protocol (Hooijmans et al., 2014a). The keywords, inclusion and exclusion criteria were planned to select publications reporting studies aiming to stimulate or inhibit the PFC, or any of its sub-divisions, before the FST in rats. Every publication included in the analysis was quali - and quantitatively evaluated to create a synthesis of the literature able to provide a robust evidence on the role of PFC on antidepressant-like effects.

## 2. Methods

### 2.1. Systematic review

The protocol was developed based on the format of the platform *Systematic Review Center for Laboratory animal Experimentation* (SYRCLE). The full text of the protocol including search strategy, screening criteria and methods for the analysis was deposited at the platform *Open Science Framework* (OSF) (Domingues et al. 2018, https://osf.io/3wh6r/), i.e., decisions about search and analytics were taken beforehand. Briefly, the search in Medline was performed in 12/12/2019 using the advanced search of the platform PubMed (http://www.ncbi.nlm.nih.gov/pubmed) with Boolean operators, and the keywords were defined according to previous protocols (Ramos-Hryb et al., 2018), as well as the keywords of articles in the research field (Table 1). Predefined inclusion and exclusion criteria were applied in two rounds of screening to obtain relevant publications as follows: 1) analysis of the titles and abstracts and 2) full texts. Reviews, systematic reviews, and meta-analysis were excluded. Original articles were included in the review when published from 1977 until the day of the search since the FST protocol was first published in that year (Porsolt et al., 1977). Due to the previous evidence showing a role of the frontal cortex in the behaviour of rats submitted to the FST (Sartim et al., 2016; Slattery et al., 2011), only studies performed in rats were included independent of the experimental design. The interventions of interest were excitation or inhibition of the PFC or its subregions using local injections, electric stimulation or other local brain manipulation techniques (i.e. DBS). Interventions were classified as excitatory or inhibitory or unknown according to the definitions of the authors of each article. Subregions of the mPFC were classified as infralimbic cortex (IL), prelimbic cortex (PL), anterior cingulate cortex (ACg), medial prefrontal cortex (mPFC) or ventromedial prefrontal cortex (vmPFC), according to the definitions reported by the authors of each article. The primary outcome was the immobility of rats in the FST reported as time, frequency, scores. Studies using rats with comorbidities or receiving co-treatment or not presenting appropriated population, interventions or outcomes were excluded. Quality assessment of the included publications was performed by using the table adapted from Risk of bias tool of SYRCLE (Hooijmans et al., 2014b).

**Table 1.**
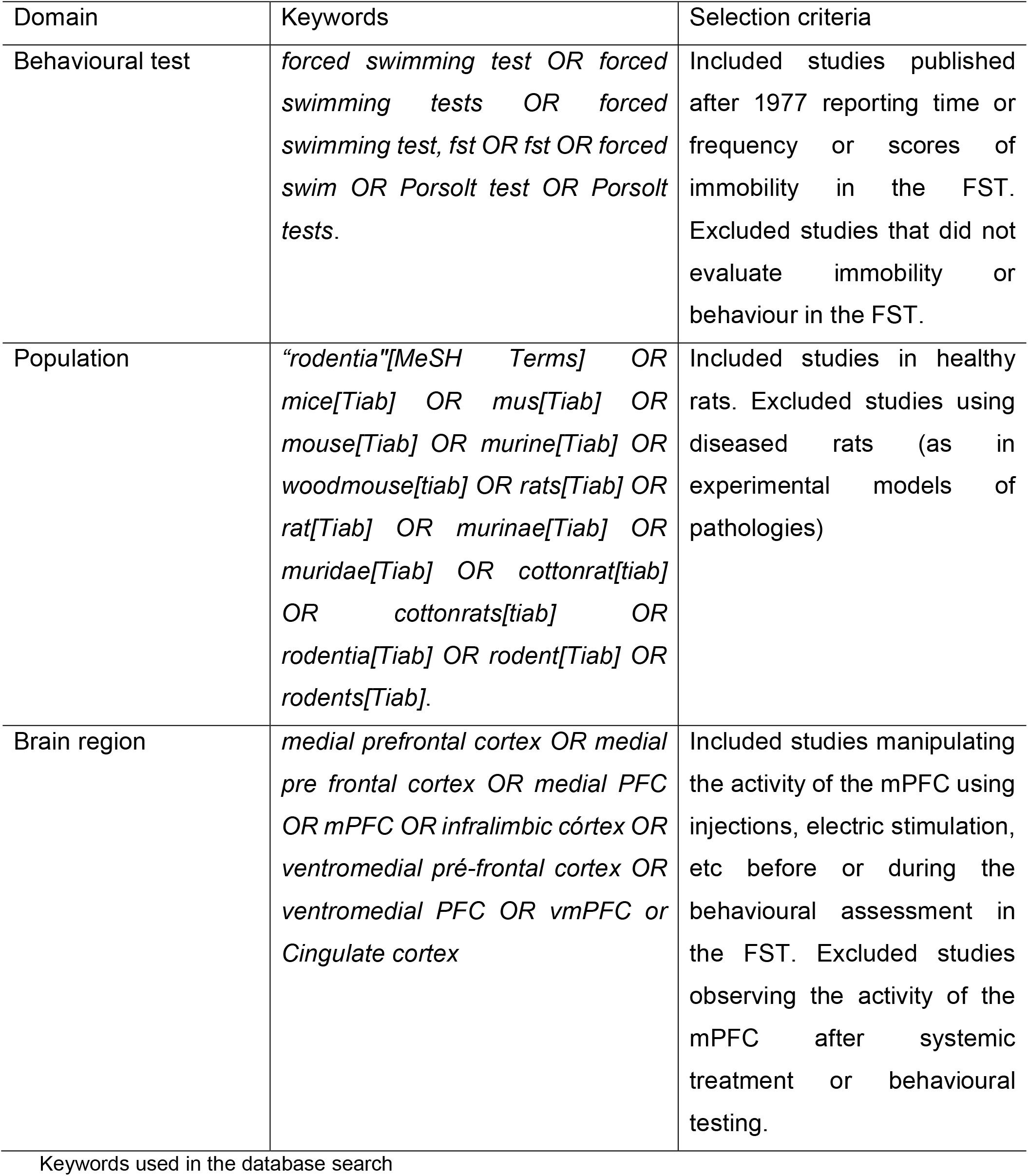
Search strategy and selection of publications:

### 2.2. Meta-analysis

For the meta-analysis (according to Domingues et al. 2018) the following parameters were extracted by KD (double-checked by FFM) from the included publications: mean, standard error, standard deviation of the primary outcome, sample size (number of animals per group; experimental or control). When available, data were extracted directly from the text or the graphical figures with a measurement ruler tool from Adobe Acrobat Reader®. When sample sizes were not clearly reported, for example, reporting only a range of the number of animals across different groups, the median of the sample size rounded down without decimals was used.

### 2.3. Analysis of the overall data

Data were meta-analyzed by random-effects model using a custom script (available in: https://osf.io/3wh6r/) written in R (http://www.r-project.org/) with the statistical package “Metafor” (free use and access, download available in http://www.metafor-project.org). Since diverse manipulations and anatomical regions were found in our systematic review could result in antagonic resuls, that is, the effect could either be reduction of increase in immobility, we chose this model of analysis because it permits two tails.

### 2.4. Effect sizes calculation

In brief, The calculation of effect sizes was performed using Hedges’ g (Hedges, 1981), given by Equation (1):

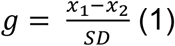

Being, *x*_1_ = mean of the experimental group; *x*_2_ = mean of the control group; SD = combined standard deviation, which is given by Equation (2):

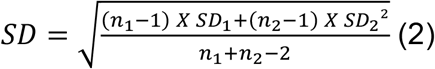

Being, n_1_ = number of animals in the experimental group; SD_1_^2^ = Standard deviation of the experimental group squared; n_2_ = number of animals in the control group; SD_2_^2^ = standard deviation of the control group squared. It is important to note that the equations were organized in a way so that reductions in immobility in the FST, in other words, the “antidepressant-like effect”, on experimental groups would yield negative effect sizes (experimental minus controls). This adaptation was done to improve data presentation since negative numbers will better imply the desired effect of the treatment (i.e. reduction of immobility). Concordantly, increased immobility would give positive g values interpreted as “stressor-like effect”.

The magnitude of effect sizes were defined as “very small” (0.01-0.2); “small” (0.2-0.5); “medium” (0.5-0.8); “large” (0.8-1.2); “very large” (1.2-2); “huge” (>2) (Sawilowsky (2009). The estimated effect size was considered “statistically significant” when 95% confidence interval (95% CI) spare the null effect and “inconclusive” when 95% CI overlap the null (Nakagawa and Cuthill (2007)

To further investigate the contribution of subgroups to the combined effect size of the overall model, a stratified meta-analysis was performed for each type of intervention (activation, inhibition, unknown) on each of the different sites, classified according to the nomenclature used by the author of the original study: IL, PL, aCg, mPFC (which includes IL + PL + acg), vmPFC (which includes IL + PL).

### 2.5. Heterogeneity analysis

In these experimental settings, although some random variation is expected due to chance alone, heterogeneity may be caused by several factors of the studies due to (known or unknown) experimental design differences. Thus, we estimated the heterogeneity of the models using the I^2^ as an indicator of the percentage of the variance between studies (Higgins et al., 2003). The magnitude of heterogeneities was classified as “very low” (0-25%); low (25-50%); “moderate” (50-75%); “high” (>75%) (HIGGINS *et al*. (2003).

### 2.6. Leave-one-out analysis

This analysis was performed for all subgroups that presented high heterogeneity, however, only in one of the iterations of the IL subgroup was possible to reduce de value of heterogeneity, therefore only this result is presented.

### 2.7. Sensitivity analysis

A sensitivity analysis, using the leave-one-out method, consisted of repeating the calculations for Hedges’ g without one of the studies in each iteration. This way, it was possible to assess the influence that each study had on both effect size and heterogeneity on the overall model

### 2.8. Publication bias analysis

The publication bias was assessed using the Trim and Fill analysis.

A symmetric funnel with no studies missing reveals the total absence of publication bias regarding this type of studies.

## 3. Results

### 3.1. Systematic review

The search in Pubmed retrieved 217 publications from which 181 were excluded from the analysis after two rounds of screening (Figure 2). Reasons for exclusion of publications were inappropriate population (n=119; 91 mice, 2 guinea pig, 1 human, 1 vole rat, 24 rats with comorbidity), intervention (n=107; 45 systemic treatment and 62 brain sites different of mPFC or subregions), outcome (n=6 without behavioural scoring) or type of publication (n=2 reviews). The 39 publications included in the analysis reported 65 different studies in 1103 adult, male rats of different strains (Table 2). The list of references with complete bibliographic information may be found at (https://osf.io/wfsv6/). None of the studies included in the review reported either sample size calculations or randomization to group allocation while about a third of them reported blindness for outcome assessment (n=14). About half of the studies did not mention randomization or blindness (n=19). Few studies reported both randomization and blindness (n=04) while one article declared no-blind assessment of the outcome (Table 2).

**Table 2:**
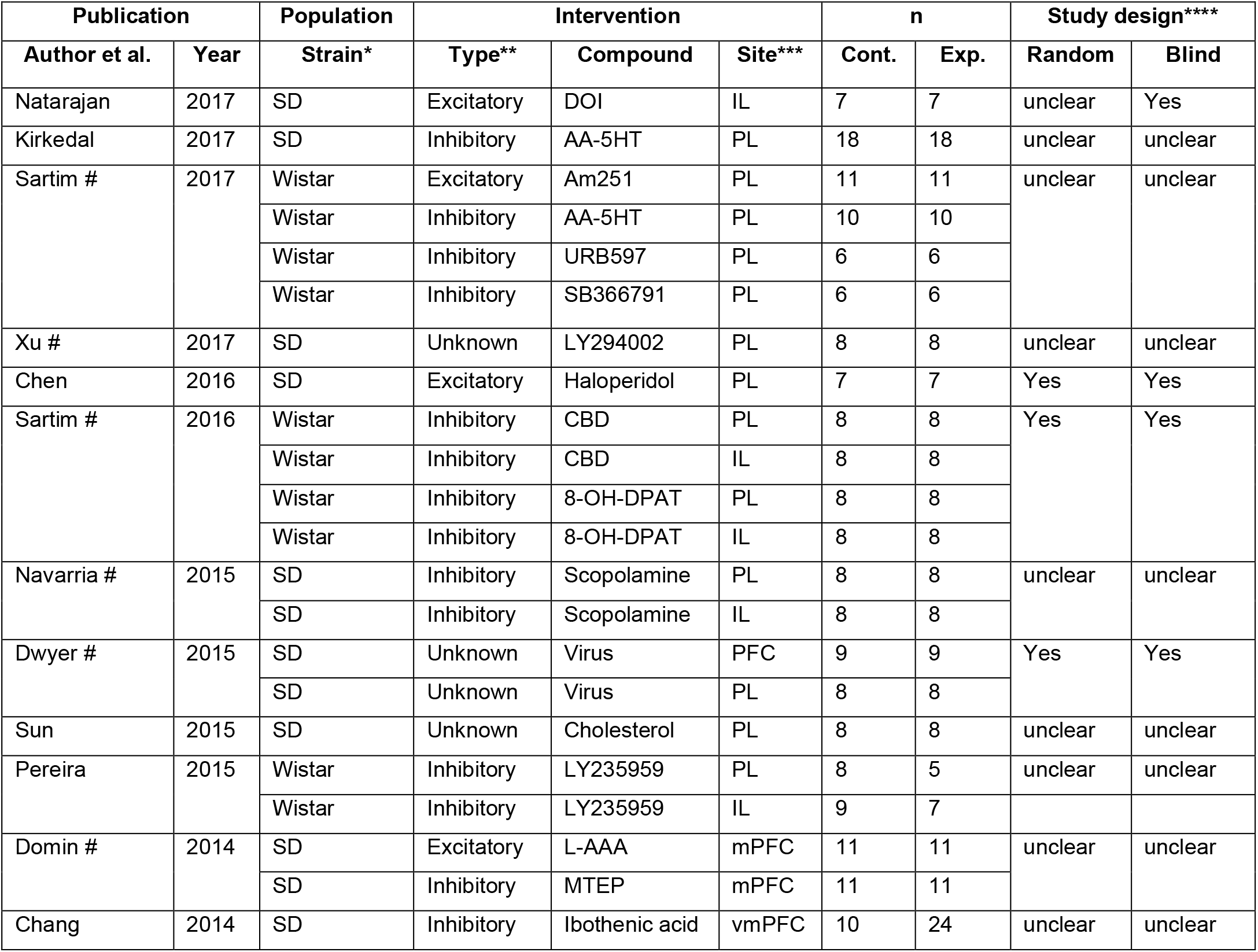
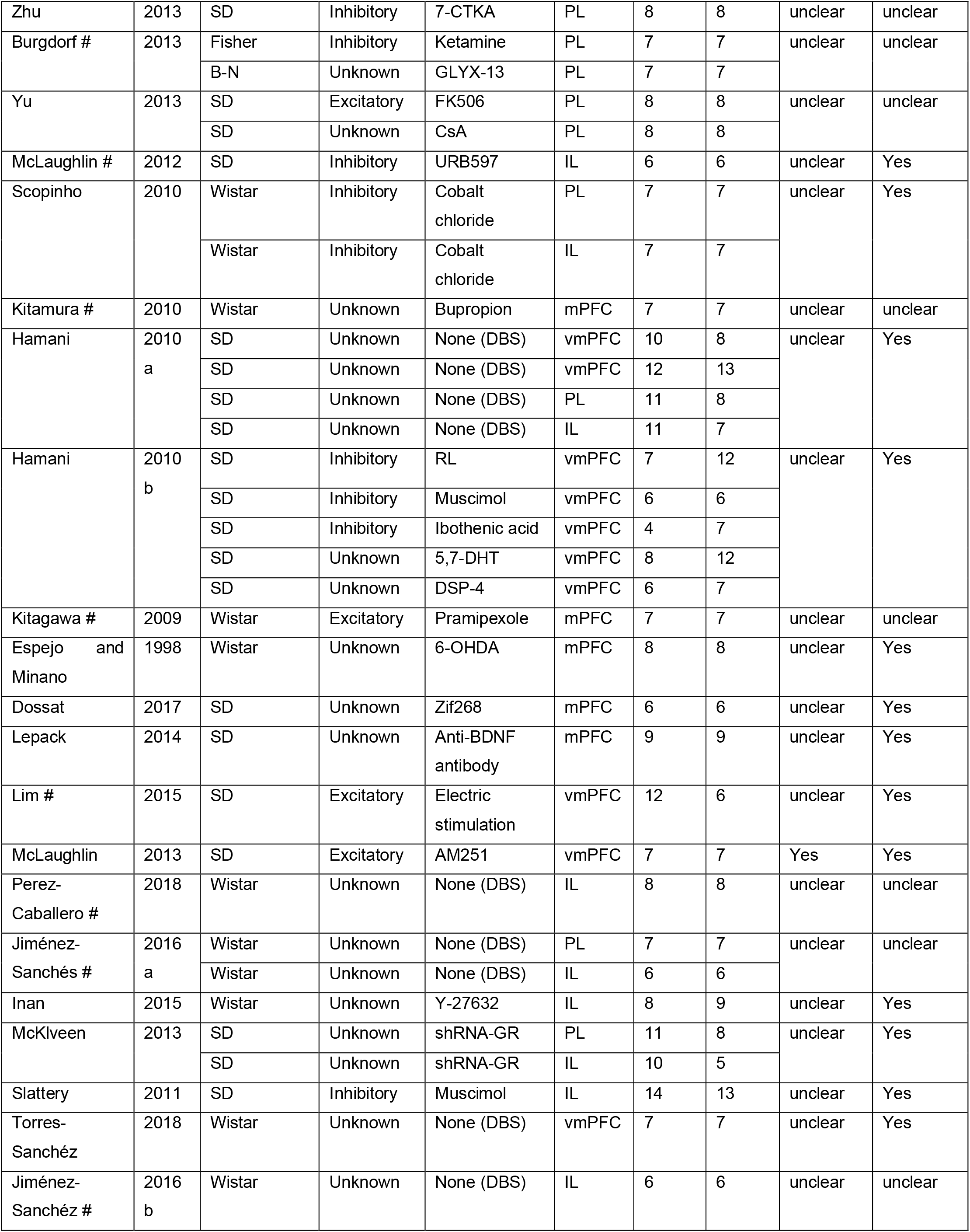
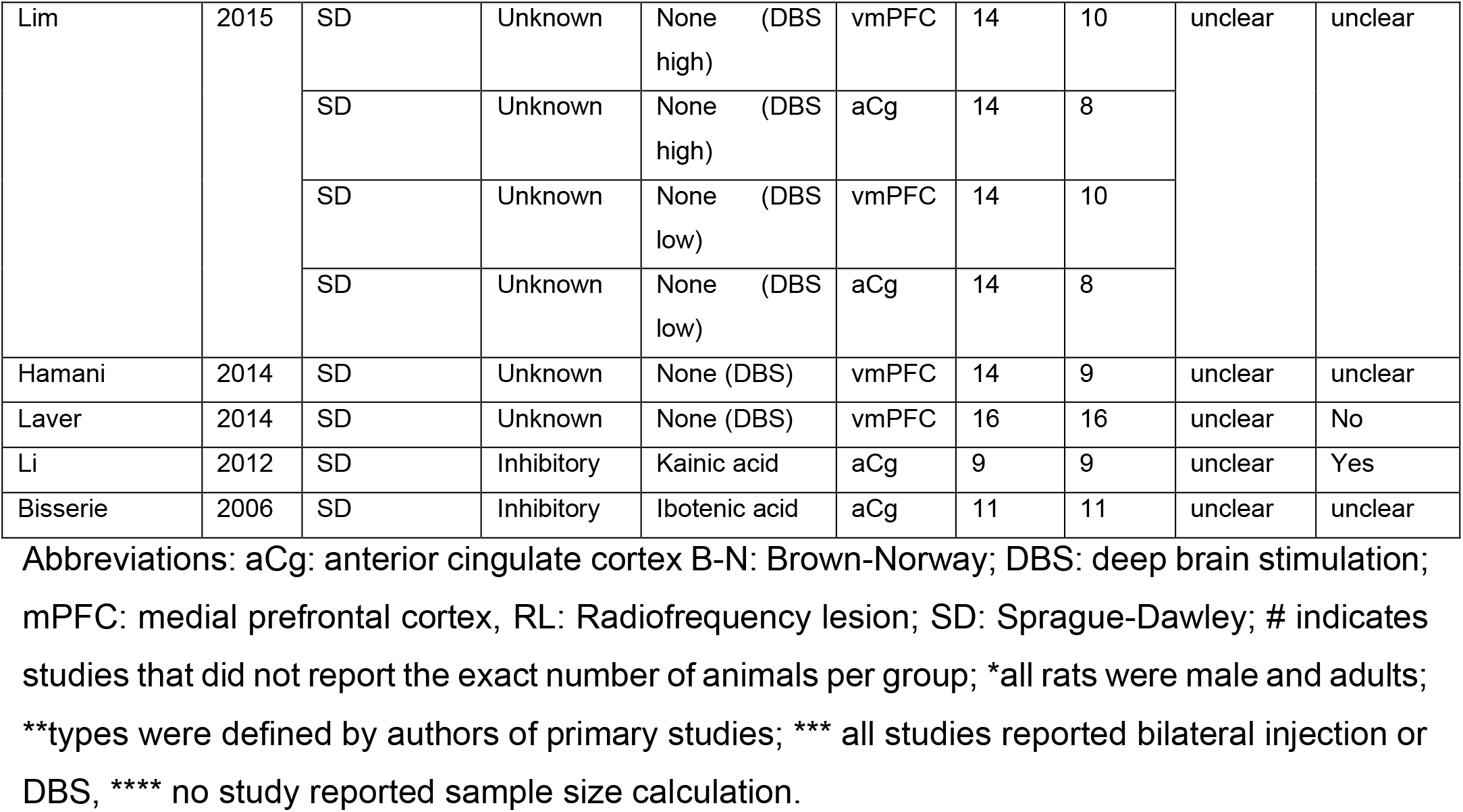

**Legend for Figure 2.**
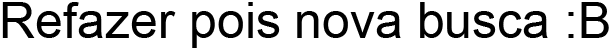
Flowchart of the systematic review and meta-analysis.

### 3.2. Meta-analysis

Overall effect size (k = 63 studies; Figure 3) was negative, i.e., in favor of the intervention, medium and significant with high heterogeneity (Hedges’ g = -0.7656, 95% CI: -1.1, -0.43; p < 0.0001; I^2^ = 84,03%). To estimate the publication bias, a funnel plot and Trim-and-fill analysis were performed. Trim-and-fill analysis suggested 12 studies missing in favor of controls/increased immobility in the FST (Figure 4), suggesting modest publication bias. The adjusted effect size was smaller than non-adjusted but still in favour of the intervention and statistically significant (Hedges’ g =-0.3964; 95% CI: -0.7577, -0.0352, p < 0.0315). The heterogeneity estimated by the adjusted model is higher than the original model (I^2^ =87.41%). In the stratified meta-analysis, heterogeneity remained high (I^2^ > 75%) for most of the subgroups, except for “no heterogeneity” for the subgroups with a single study (“activation IL”, “inhibition mPFC”); very-low for “unknown aCg”; low for “inhibition PL” medium heterogeneity for “unknown vmPFC”.

**Legend for Figure 3.**
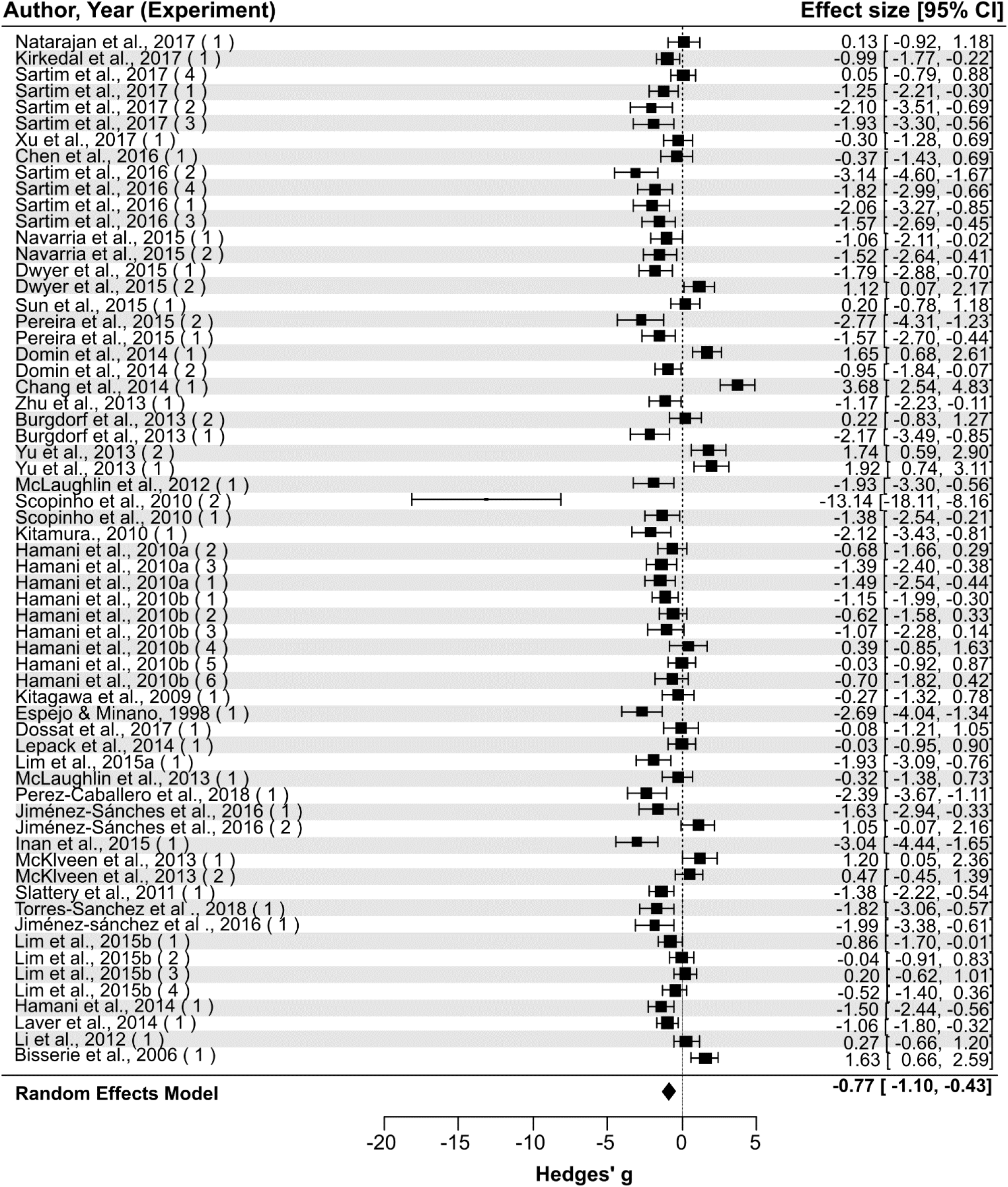
Forest Plot and results of the metanalysis with random-effects model. Showing the effects observed for each study (black squares) with its respective 95% confidence intervals (horizontal lines, CI) and the effect of the total metanalysis (diamond). Sample sizes may be found in table 2. Symbols located at the left side of the dotted line (representing the null effect) indicate interventions reducing the immobility time in the FST (antidepressant-like effect). In contrast, those at the right side indicate intervention increasing immobility time (stressor-like effect).

**Figure 4.**
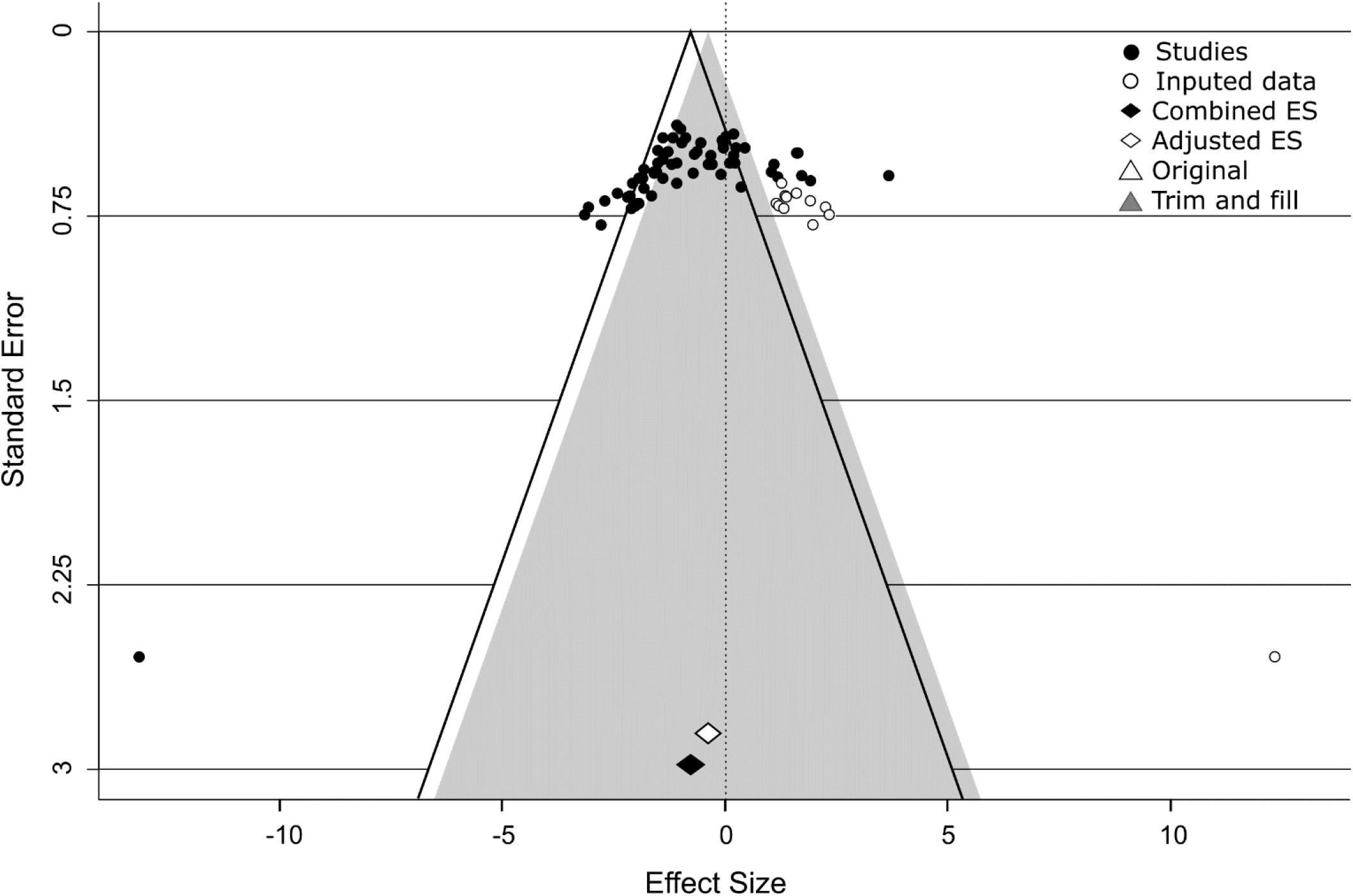
Funnel plot and Trim-and-fill analysis. Effect sizes (ES) calculated from the 63 studies (black dots) included in the original meta-analysis plotted against their errors yielded a slightly asymmetric funnel (black lines). Trim-and-fill analysis indicated 12 studies missing (blank circles) from original data in order to create symmetry in the funnel plot (shaded triangle). Non-adjusted combined effect size (black diamond) was towards the intervention side and statistically significant. Adjusted combined effect size (white diamond) was towards the intervention side, statistically significant but smaller than non-adjusted combined effect size.

For the subgroup “activation mPFC” (Figure 5 a; k = 2 studies) there was a medium, non-significant increase of immobility with high heterogeneity (Hedges’ g = 0.6982; 95% CI: -1.18, 2.58; p=0.4664; I^2^ = 85.52%). For the subgroup “inhibition mPFC” (Figure 5 b, k = 1 study), there was a large, significant reduction of immobility (Hedges’ g = -0.9531; 95% CI: -1.84, -0.07; p=0.0342) with no heterogeneity since there was a single study. The subgroup of the “unknown mPFC” (Figure 5 c; k = 4 studies) presented a very-large, non-significant reduction of immobility with high heterogeneity (Hedges’ g = -1.17092; 95% CI: -2.52, 0.17; p=0.0881; I^2^ = 81.28%).

**Figure 5.**
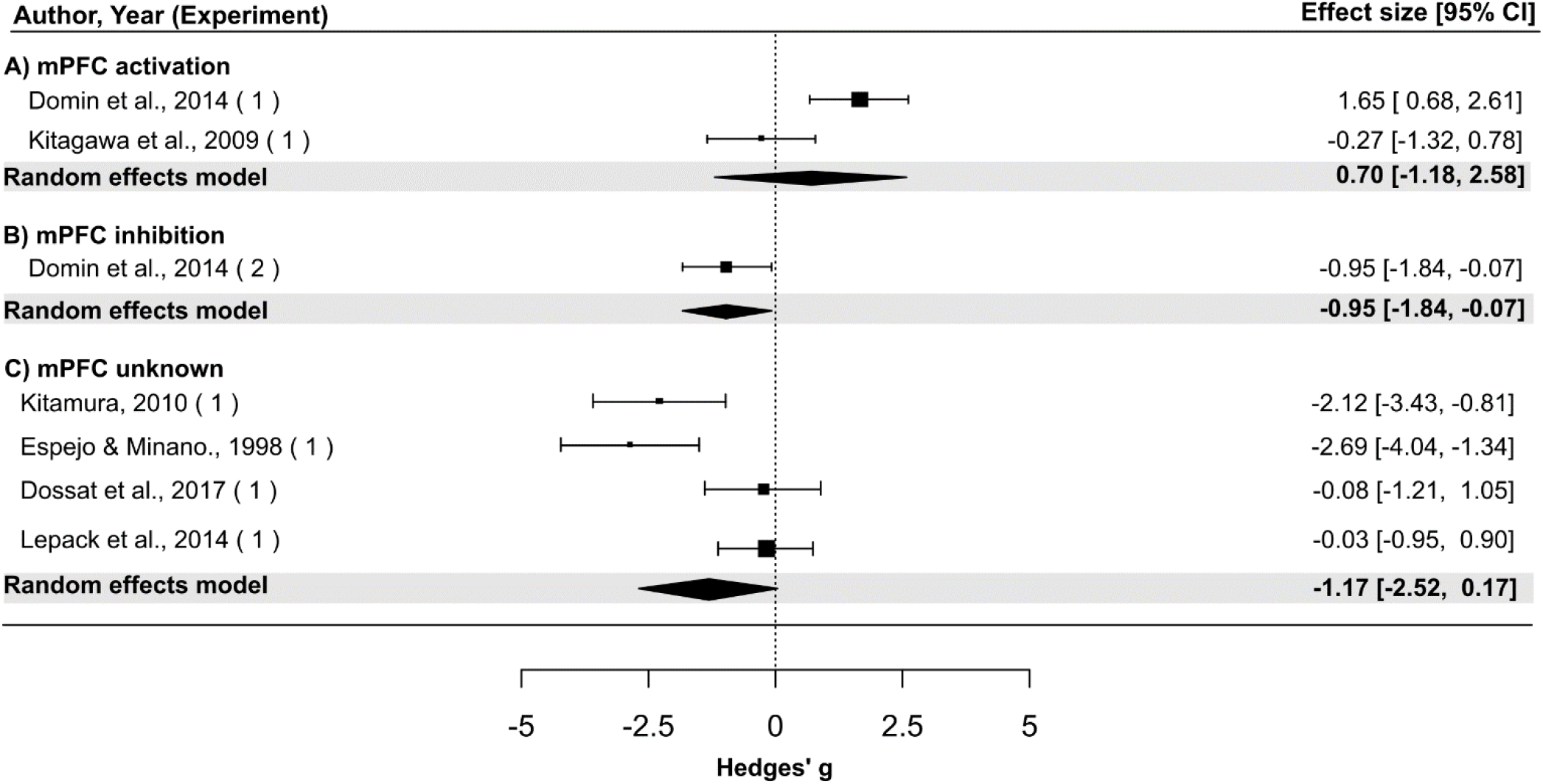
Forest Plot and metanalysis result of the subgroup mPFC (medial prefrontal cortex) with the random-effects model. Showing the effects observed for each study (black squares) with their 95% confidence intervals (horizontal lines, CI) and metanalysis stratified by interventions (activation, inhibition, unknown, black diamonds). Sample sizes may be found in table 2. Symbols located at the left side of the dotted line (representing the null effect) indicate interventions reducing the immobility time in the FST (antidepressant-like effect). In contrast, those at the right side indicate intervention increasing immobility time (stressor-like effect).

The subgroup “activation vmPFC” (Figure 6 a, k = 2 studies) presented a large, non-significant reduction of immobility with high heterogeneity (Hedges’ g = -1.1057; 95% CI: -2.68, 0.47; p=0.1682; I^2^ = 75.06%). The subgroup “inhibition vmPFC” (Figure 6 b, k = 4 studies), there was a medium, non-significant increase of immobility with high heterogeneity (Hedges’ g = 0.5907; 95% CI: -1.51, 2.70; p=0.5824; I^2^ = 92.82%). For the subgroup of the “unknown vmPFC” (Figure 6 c, k = 9 studies), there was a large, significant reduction of immobility with medium heterogeneity (Hedges’ g = -0.8859; 95% CI: -1.32, -0.45; p < 0.0001; I^2^ = 50.23%).

**Figure 6.**
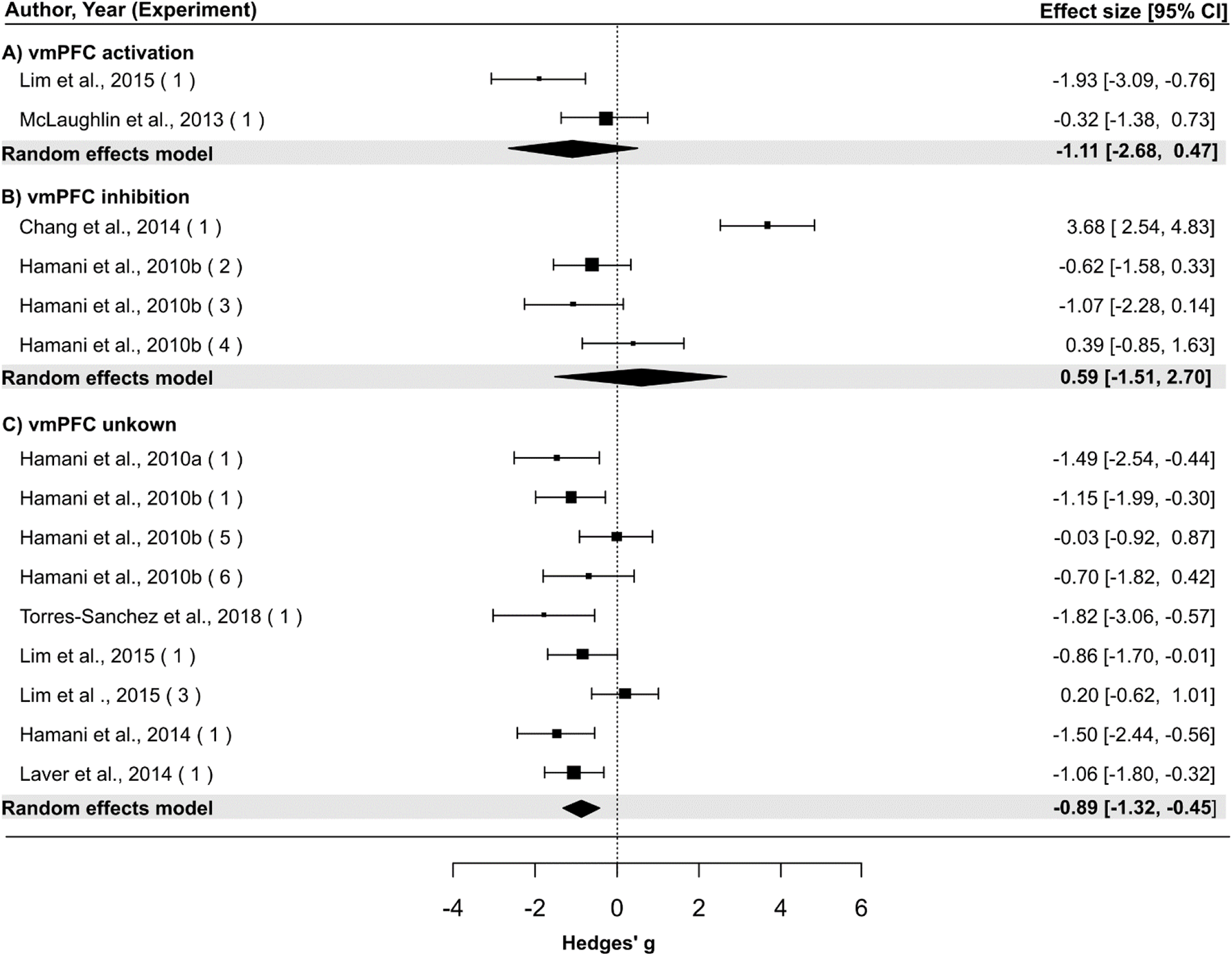
Forest Plot and metanalysis result of the subgroup vmPFC (ventral medial prefrontal cortex) with random-effects model. Showing the effects observed for each study (black squares) with their 95% confidence intervals (horizontal lines, CI) and metanalysis stratified by interventions (activation, inhibition, unknown, black diamonds). Sample sizes may be found in table 2. Symbols located at the left side of the dotted line (representing the null effect) indicate interventions reducing the immobility time in the FST (antidepressant-like effect). In contrast, those at the right side indicate intervention increasing immobility time (stressor-like effect).

There were no cases in the subgroup “activation aCg” (Figure 7 a). The subgroup “inhibition aCg” (Figure 7 b; k = 2 studies) presented a large, non-significant increase of immobility with high heterogeneity (Hedges’ g = 0.9445; 95% CI: -0.38, 2.27; p=0.1636; I^2^ = 74.63%). For the subgroup “unknown aCg” (Figure 7 c, k = 2 studies), there was a small, non-significant reduction of immobility with null heterogeneity (Hedges’ g = -0.2774; 95% CI: -0.90, 0.34; p=0.3797; I^2^ = 0%).

**Figure 7.**
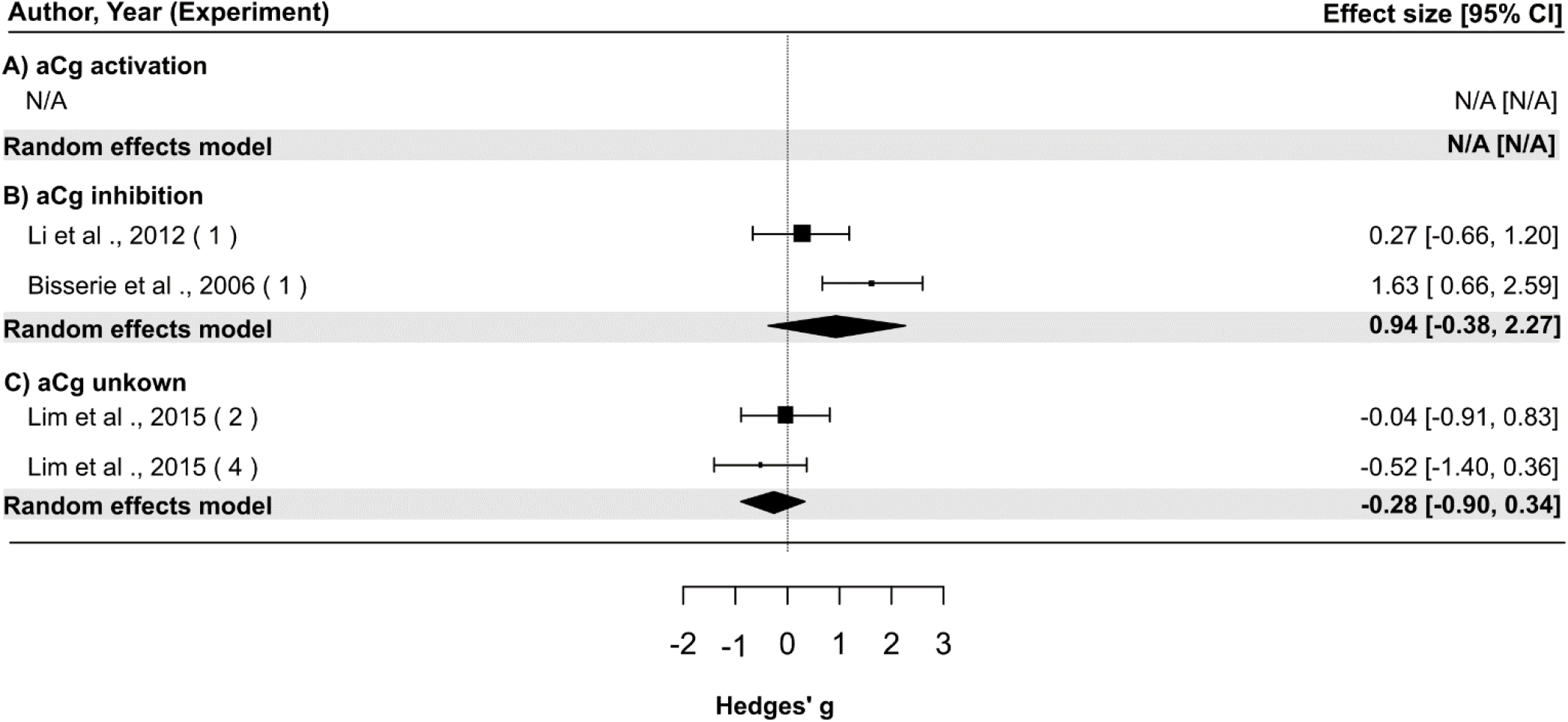
Forest Plot and metanalysis result of the subgroup aCg (anterior cingulate cortex) with random-effects model. Showing the effects observed for each study (black squares) with their 95% confidence intervals (horizontal lines, CI) and metanalysis stratified by interventions (activation, inhibition, unknown, black diamonds). Sample sizes may be found in table 2. Symbols located at the left side of the dotted line (representing the null effect) indicate interventions reducing the immobility time in the FST (antidepressant-like effect). In contrast, those at the right side indicate intervention increasing immobility time (stressor-like effect).

For the subgroup “activation PL” (Figure 8 a; k = 3 studies), there was a medium, non-significant increase of immobility with high heterogeneity (Hedges’ g = 0.4404; 95% CI: -0.78, 1.76, p=0.4786; I^2^ = 77.12%). For the subgroup “inhibition PL” (Figure 8 b; k = 11 studies), there was a very large, significant reduction of immobility with low heterogeneity (Hedges’ g = -1.3091; 95% CI: - 1.68, -0.93; p < 0.0001; I^2^ = 22.67%). For the subgroup “unknown PL” (Figure 8 c; k = 9 studies), there was a very small, non-significant reduction of immobility with high heterogeneity (Hedges’ g = - 0.0883; 95% CI: -0.99, 0.82, p=0.8484; I^2^ = 84.68%).

**Figure 8.**
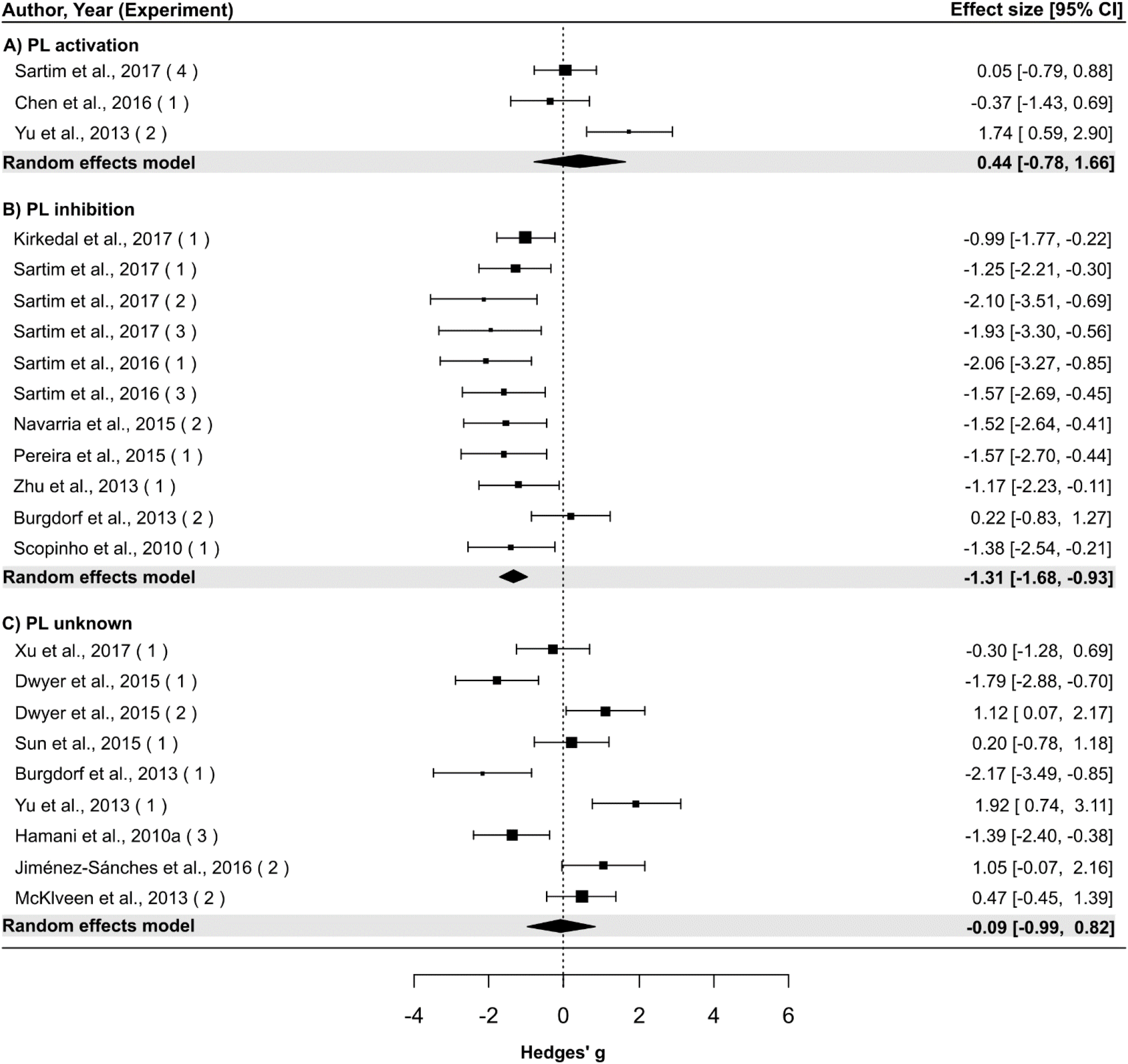
Forest Plot and metanalysis result of the subgroup PL (pre limbic cortex) with random-effects model. Showing the effects observed for each study (black squares) with their 95% confidence intervals (horizontal lines, CI) and metanalysis stratified by interventions (activation, inhibition, unknown, black diamonds). Sample sizes may be found in table 2. Symbols located at the left side of the dotted line (representing the null effect) indicate interventions reducing the immobility time in the FST (antidepressant-like effect). In contrast, those at the right side indicate intervention increasing immobility time (stressor-like effect).

The subgroup “activation IL” (Figure 9 a, k = 1 study) presented a small, no-significant increase of immobility (Hedges’ g = 0.1319; 95% CI: -0.92, 1.18; p = 0.8067) with null heterogeneity since there was a single study in the subgroup. For the subgroup “inhibition IL” (Figure 9 b, k = 7 studies), there was a huge and significant reduction of immobility with high heterogeneity (Hedges’ g = -2.8942; 95% CI: -4.87, - 0.92; p = 0.0040; I^2^ = 93.49%). Leave-one-out analysis showed heterogeneity remained high with removal of individual studies, except by excluding Scopinho et al. (2010) (Figure 10). Removal of Scopinho et al. (2010) reduced heterogeneity to low levels and CES to very large (I^2^ = 32.8842%; Hedges’ g = -1.8484; 95% CI: -2.44, -1.25; p < 0.000001; Figure 10). For the subgroup “unknown IL” (Figure 9 c, k = 6 studies), there was a large, significant reduction of immobility with high heterogeneity (Hedges’ g = -1.3856; 95% CI: -2.60, -0.17, p = 0.0256) (I^2^ = 82,85%).

**Figure 9.**
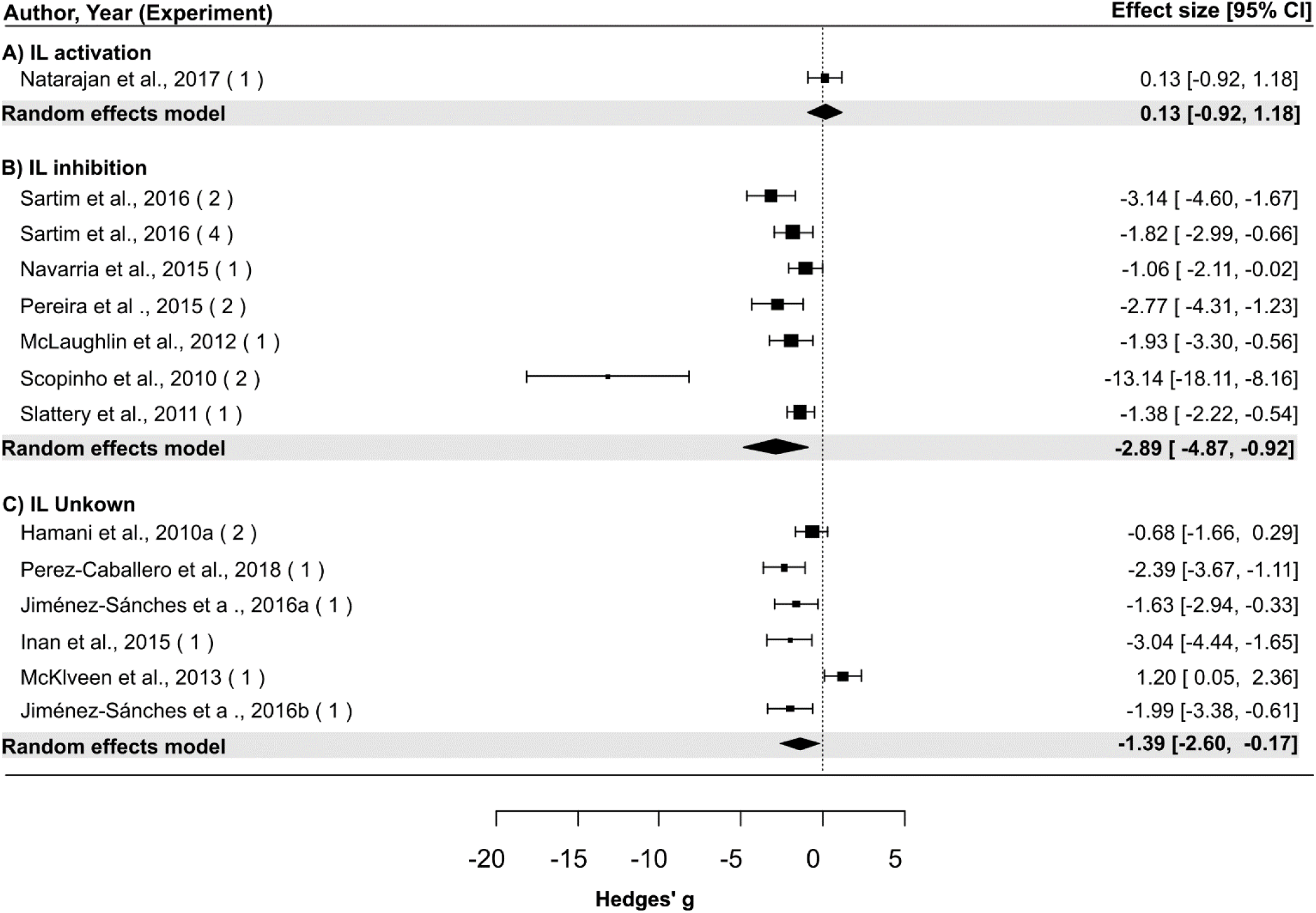
Forest Plot and metanalysis result of the subgroup IL (infra limbic cortex) with random-effects model. Showing the effects observed for each study (black squares) with their 95% confidence intervals (horizontal lines, CI) and metanalysis stratified by interventions (activation, inhibition, unknown, black diamonds). Sample sizes may be found in table 2. Symbols located at the left side of the dotted line (representing the null effect) indicate interventions reducing the immobility time in the FST (antidepressant-like effect). In contrast, those at the right side indicate intervention increasing immobility time (stressor-like effect).

**Figure 10.**
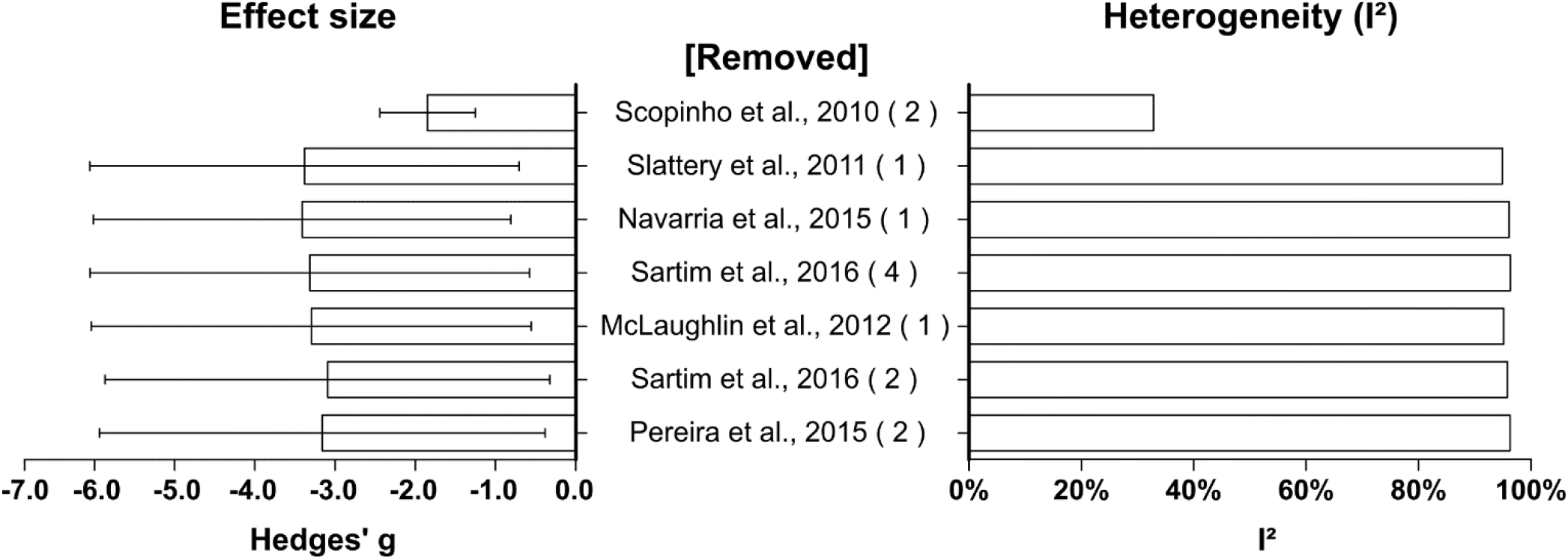
Sensitivity analysis “*leave-one-out”* of the subgroup “Infralimbic inhibition”. The bars represent the combined effect size (Hedges’ g, left side) or heterogeneity (I^2^, right side) calculated in the absence of the study mentioned in the respective line in the middle column.

## 4. Discussion

This is the first quantitative review on the causal relationship between the intervention in the excitability of the mPFC and behaviour of rats in the FST. A systematic review and meta-analysis were performed to determine the direction, magnitude, significance, heterogeneity of effect sizes and the risk of biases in the field of research. The present data indicates that the combined effect size for the whole model was medium-sized, negative (i.e. antidepressant-like effect) and significant with high heterogeneity. The combined effect size, differently interpreted according to the research field, varied in magnitude when corrected by publication bias or stratified within types and sites of interventions. However, the lack of a consensus in the research field, very small or small effects may be considered less relevant than the medium, large or huge ones (Bezeau and Graves, 2001).

Combination of different types of bias may contribute to inflate the estimated effect size of an intervention distorting the appraisal of the efficacy (SENA et al, 2010; MACLEOD et al., 2015; RAMOS-HRYB et al 2018). Trim-and-fill analysis of the whole data set (k=63) suggested 12 studies missing from the original model, indicating a possible publication bias towards the intervention. In fact, the adjusted effect size had a smaller magnitude and higher heterogeneity than the non-adjusted one, but still statistically significant. Risks of other types of bias (e.g. attrition and reporting, HOOIJMANS et al, 2014) were not appraised. Risk of selection, performance, and detection biases (HOOIJMANS et al, 2014) may not be discarded once that is unclear how randomization and blinding were used in most the publications, contributing to inflating the estimated effect size.

Conversely, subgroups of studies reporting effect sizes against the intervention or with small magnitude may deflate the combined effect size spuriously (LINO DE OLIVEIRA et al., 2020). Indeed, stratification of data by type of intervention and subregion of the mPFC yield antidepressant-like, statistically significant effects considered large, very large or even huge with the inhibition of the IL, PL or mPFC. In contrast, interventions that potentially increased neural activity of a given subregion of the PFC showed non-significant and predominantly positive (stressor-like effect) effect sizes, except for vmPFC in which a negative, albeit the non-significant effect was observed. Interestingly, interventions with unknown effects on the neural activity, had neglectable effects when applied to PL and aCg while prominent antidepressant-like effect when applied to IL, vmPFC or mPFC. However, none of the effects were statistically significant.

The huge magnitude of the effect sizes of inhibiting the IL suggested a higher contribution of this region to the control of rat immobility in the FST when compared to the large and very large magnitudes of inhibiting mPFC or PL, respectively. Further leave-one-out analysis of data suggests that the compound used to inhibit the IL may play a role in the magnitude of the effect size. Exclusion of Scopinho *et al. (2010*), a study using cobalt chloride to inhibit the IL with an effect size 5 times larger than the combined one, reduced the effect size and of the IL inhibition to values closer to, but still higher than, inhibition of PL. Cobalt chloride may cause more substantial effects since it is expected a total inhibition of a region, with a fast effect and recovery when compared to neuropharmacological inhibitions (Kretz, 1984). The hierarchical classification of the effect sizes, based on their magnitudes, could indicate a decrescent order of importance to IL, PL and mPFC in the antidepressant-like effects in rats in the FST. These last speculations require further research.

The leave-one-out analysis also showed that the exclusion of an outlier study greatly decreased the heterogeneity of the subgroup “IL inhibition”. In the remaining subgroups, null, very low, low or medium levels of heterogeneity varied with different types of intervention, subregion and number of studies examined. Null heterogeneity was observed when a single study was examined (k= 1) impairing a meta-analysis and heterogeneity calculations such as “activation IL” “inhibition mPFC” but also in the subgroup “unknown ACg” with k=2 studies. In the meta-analysis for “unknown vmPFC” (k= 9 studies) heterogeneity was considered medium. The heterogeneity was low in the subgroup “inhibition of PL” (k=11 studies). Subgroups vmPFC and mPFC presented high heterogeneity (I^2^ > 75%) similar to the overall heterogeneity. Magnitudes of heterogeneity represent the total amount of explained and unexplained variability due to known or unknown differences within experimental settings, here estimated as the percentage of variance between studies (Higgins et al., 2003). Fall in the levels of heterogeneity from the overall to subgroups, as observed for inhibition of IL, PL and mPFC or DBS in the vmPFC, suggest that type of intervention explained most of the variability. In contrast, high heterogeneity in the subgroups vmPFC and mPFC indicate an important contribution of unknowns to the observed heterogeneity. Different substances injected into the discrete cortical sites may explain the residual variability, but the myriad of compounds precluded an analysis of the subgroup.

Pharmacological interventions were compounds with diverse mechanisms of action, their inhibitory or excitatory actions on neural activity of a targeted brain region was presumed by authors interpretation since no study reported direct measures of it. Except for the ketamine (Burgdorf et al. (2013), all other presumably inhibitory compounds of the PL or IL had an antidepressant-like effect (agonists of the receptors GABA_A_ (Muscimol: Slattery et al. (2011)), cannabinoids CB_1_ and CB_2_ (CBD: Sartim et al. (2016)), serotonergic 5-HT_1A_ (8-OH-DPAT: Sartim et al. (2016)); and antagonists of glutamate NMDA, mGlu_5_ receptors (LY235959: Pereira et al. (2015); MTEP: Domin et al. (2014); 7-CTKA: Zhu et al. (2013); Ketamine: Burgdorf et al. (2013)),. In contrast, putatively excitatory compounds such as agonists of serotoninergic 5-HT_2A/C_ receptors (DOI: Natarajan et al. (2017)), dopaminergic D_2_, D_3_ and D_4_ receptors (pramipexole: Kitagawa et al. (2009)), antagonist of dopaminergic D_2_ receptors (haloperidol: Chen et al. (2016)) and cannabinoid CB_1_ (AM251: Sartim et al. (2016)) had neglectable effects on immobility time when injected into PL or IL. These data indicated an equally prominent role for the receptors GABA_A_, CB_1_, 5-HT_1A_ or NMDA and a less critical to the D_2_, D_3_ and D_4_ dopaminergic and 5-HT_2A/C_ of IL or PL on the control of the immobility behaviour in the FST. However, no empirical data exist to support the last assertions.

Studies in which DBS was applied towards any cortical site were assigned to the subgroup “unknown”, in regard of the consequence to the neuronal activity of the targeted brain region. DBS and other interventions with unknown effects on cortical excitability, assigned to the subgroup “unknown”, yielded a large, significant, antidepressant-like effect in the FST. The authors of the studies did not measure neural activity during or after the intervention nor speculated about it (Hamani et al., 2014; Hamani et al., 2010; Jiménez-Sánchez et al., 2015; Jiménez-Sánchez et al., 2016; Laver et al., 2014; Lim et al., 2015; Perez-Caballero et al., 2018; Torres-Sanchez et al., 2018). Similar to temporary (CoCl_2:_ Scopinho et al. (2010); Ibotenic Acid: Chang et al. (2014)) or permanent inactivation (Kainic acid: Li et al. (2012); radiofrequency lesions most of the studies applying DBS towards the IL or PL yielded antidepressant-like effect in the FST indicating a possible inhibition of neural activity in these brain structures. This last supposition was likely to be the more prudent, considering that the mechanisms underlying DBS effects on nervous tissue are poorly understood, as well as its ramifications to the overall neuronal function of the target region or its afferent/efferent regions (Chiken and Nambu, 2016; McIntyre et al., 2004).

The heterogeneity in the overall meta-effect may also arise from the variability in the subdivisions of the mPFC that are targeted by the various interventions, or simply from the way the neuroanatomy of the region is approached in the different studies. In rats, the subregions of the mPFC have distinct cytoarchitectural and hodological characteristics (Burgos-Robles et al., 2013; Hoover and Vertes, 2011; Hurley et al., 1991; Vertes, 2004). Thus, differences in function regarding behavioural control are to be expected. Studies that utilized inhibition protocols that encompassed large regions of the PFC yielded inconclusive results (vmPFC) or small effect sizes (mPFC). These inconclusive results may occur due to the lack of spatial resolution of the intervention site since inhibiting multiple subregions at the same time could lead to antagonistic behavioural outcomes.

In summary, the present systematic review and meta-analysis indicate that the inhibition of either IL or PL was necessary and sufficient to produce an antidepressant-like effect in male rats in the FST. Moreover, data from the literature suggest that the inhibition of the IL or PL regions mediated by the combination of different systems may be a mechanism underlying the effect of antidepressant drugs.

## Acknowledgements

Karolina Domingues was supported by Conselho Nacional de Desenvolvimento Científico e Tecnológico (140007/2016-4), Brazil. Fernando F.F.M was supported by Post-doctoral fellowship grant #2018/25857-5, São Paulo Research Foundation (FAPESP), Brazil. This study was financed in part by the Coordenação de Aperfeiçoamento de Pessoal de Nível Superior – Brasil (CAPES) – Finance Code 001”.

